# Chromosome-level genome assembly of Norwegian wild alpine reindeer (*Rangifer tarandus tarandus*)

**DOI:** 10.1101/2025.05.15.652595

**Authors:** Ole K. Tørresen, Ave Tooming-Klunderud, Morten Skage, Anne Eline Streitlien, Olav Strand, Christer M. Rolandsen, Giada Ferrari, José Cerca, Atle Mysterud, Kjetill S. Jakobsen

## Abstract

We describe a chromosome-level genome assembly from a wild alpine reindeer individual (*Rangifer tarandus tarandus)* from the Rondane area in Southern Norway. The assembly is resolved into two pseudo-haplotypes: hap 1 spanning 3,081 megabases and hap 2 spanning 2,633 megabases. Contig N50 and scaffold N50 lengths are in the range of 31-41 Mb and 66-69 Mb, respectively. A large part of these two haplotypes (83.8% and 90.4%, respectively) are scaffolded into 34 autosomal chromosomal pseudomolecules, and in sex chromosomes X and Y for hap 1. The BUSCO completeness scores are 98.0% and 95.2%, respectively, and gene annotations of the assemblies identified 37,998 and 36,977 protein-coding genes. We also present an updated and improved genome assembly for Svalbard reindeer (*Rangifer tarandus platyrhynchus)* and a comparison with previously published genome assemblies of reindeer.

## Introduction

Reindeer is an iconic species of the northern hemisphere playing a key role in the ecosystems and also as part of human culture (Mustonen 2022). The taxonomic delimitation of the species is somehow disputed with reindeer in Eurasia (hereafter reindeer) and caribou in North America being the same species, *Rangifer tarandus*, but typically separated into a number of subspecies (Banfield 1961; Tryland and Kutz 2018). Reindeer are distributed across the northern Holarctic region, inhabiting both arctic tundra and boreal forests. Wild reindeer populations count just below 3 million adult individuals, but numbers are declining globally and they are now Red Listed as ‘Vulnerable’ (Gunn 2015). While reindeer conservation is important in itself, this species also functions as an umbrella for conservation of a wider biodiversity (Labadie et al. 2024).

Reindeer has a polygynous mating system, and are sexually dimorphic in body size with males being considerably larger than females. Reindeer is the only cervid where females also possess antlers. Reindeer have a series of unique biological features enabling survival in harsh arctic environments. Reindeer are adapted to winter darkness and can see wavelengths down into the ultraviolet range (Stokkan et al. 2013), and they have specific mutations in genes involved in the vitamin D and fat metabolism pathways (Lin et al. 2019). They have a unique ability to digest and rely on a winter diet consisting of a high proportion of lichens (Falldorf et al. 2014). Reindeer have the most extreme snow coping abilities in terms of morphology among cervids (Telfer and Kelsall 1984).

Reindeer have been semi-domesticated at least two times (Røed et al. 2008) and there is evidence for milder artificial selection relative to fully domesticated species (Wu et al. 2024). Reindeer herding has been a cornerstone to the culture of several Eurasian nomadic indigenous people (Salmi 2022). Semi-domestic reindeer have also been introduced to areas in Norway, Iceland, Greenland, Alaska, and South Georgia island to live as wild, i.e., becoming feral reindeer (Mysterud, Flagstad, and Strand 2024). In Fennoscandia, semi-domestic reindeer have replaced wild reindeer from most parts of their former range, and admixture with the semi-domestics threaten the genetic integrity of the remaining wild reindeer (Anderson et al. 2017). A key issue for reindeer conservation in Europe is to save the last wild reindeer, including mainly alpine-living reindeer (*Rangifer tarandus tarandus*) in Norway and forest-living reindeer (*Rangifer tarandus fennicus*) in Finland (Weldenegodguad et al. 2024). This is challenging due to partial mixing with semi-domestic and feral populations. Norway harbors more than ∼270,000 semi-domestic and feral populations with high levels of genetic mixing with semi-domestics, but only some 8,000 wild reindeer with a low level of genetic mixing (Kvie et al. 2019).

Climate change, fragmentation and habitat loss are well-known challenges for the conservation of reindeer (Dorber et al. 2023; Morineau et al. 2023). There are several gaps related to understanding the role of genetics critical to target conservation efforts. Fragmentation leads to smaller populations and with less gene flow to adjacent populations, and these are likely subject to increasing risk of inbreeding, drift and loss of genetic diversity (Kvie et al. 2019; Hansen et al. 2024), similar processes as in the overharvesting of the Svalbard reindeer in 17-20th centuries (Kellner et al. 2024). The issue of reindeer genetics is currently also made topical due to an outbreak of chronic wasting disease (CWD) in Norway. This led to depopulation of the Nordfjella reindeer area in Norway with culling of more than 2,000 wild reindeer (Mysterud and Rolandsen 2018), and harvest management leading to reduced population size and highly skewed sex-ratio with few males in the largest wild reindeer population in Norway (Mysterud et al. 2025).

There are several genome assemblies for reindeer (Dussex et al. 2023; Z. Li et al. 2017; Poisson et al. 2023), and genome assemblies at chromosome-level already exist (Poisson et al. 2023; Dussex et al. 2023). One reference genome stems from a one-year-old male reindeer from Sodankylä, Finland (Pokharel et al. 2023; Weldenegodguad et al. 2020), representing a semi-domestic reindeer. A high-quality reference genome is available for the Svalbard reindeer (*Rangifer tarandus platyrhynchus*) (Dussex et al. 2023). This subspecies is atypical having lived on an arctic island without predators for about 5,000 years, and they are much less social, sedentary, smaller in body size, have shorter legs, store more fat and have limited genetic variation.

Almost all of the remaining wild alpine reindeer in Europe are in Norway (Gunn 2015). Of the 24 populations managed as wild reindeer in Norway, only **four of these populations** (Snøhetta, Knutshø, Rondane, and Sølnkletten) have limited admixture with semi-domestics, and are considered vulnerable due to ongoing habitat fragmentation and a decline in performance indicators. Here, we have generated a chromosome-level, haplotype-resolved genome assembly of the wild alpine reindeer (*R. tarandus tarandus*) genome created using PacBio HiFi long-read and Hi-C sequencing data and compare it to other publicly available genomes. This haplotype-resolved genome assembly is generated as part of the Earth Biogenome Project Norway.

## Methods

### Sample acquisition and DNA extraction

The sample was from a 3.5 year old male reindeer from Jervetunga, Rondane, Norway (UTM32 533028, 687324). The sample was from a reindeer legally shot during the ordinary hunting season on August 29, 2023, and sampling from such a specimen requires no specific legal permit in Norway. The reference code for the sample is 100000423715 in the registry for deer data in Norway (‘Hjorteviltregisteret’).

DNA isolation for PacBio long-read sequencing was performed using a QIAGEN Genomic-tip 100/G gravity-flow column with 78 mg of flash-frozen heart tissue as starting material and following the manufacturer’s instructions for tissue samples. Quality check of amount, purity and integrity of isolated DNA was performed using the Qubit BR DNA quantification assay kit (Thermo Fisher), Nanodrop (Thermo Fisher), and Fragment Analyser (DNA HS 50 kb large fragment kit, Agilent Tech.).

### Library preparation and sequencing for *de novo* assembly

Approximately 8 µg of purified HMW DNA was pre-conditioned with the DNAFluid+ kit and sheared into an average fragment size of 30 kb using the Megaruptor3 (Diagenode). For library preparation, 5 µg of fragmented DNA was used following the PacBio protocol for HiFi library preparation using the SMRTbell® prep kit 3.0. The final HiFi library was size-selected with a 10 kb cut-off using a gel cassette on the BluePippin instrument (Sage Science), followed by an additional bead clean-up (PacBio) and library quantification with Qubit HS DNA quantification assay kit (Thermo Fisher). Final library size estimation was done with the Fragment Analyser kit mentioned above. Sequencing was performed by the Norwegian Sequencing Centre on the PacBio Revio instrument using the Revio™ polymerase kit and one 25M SMRT cell.

A Hi-C library was prepared using the Arima High Coverage HiC kit (Arima Genomics, Inc), following the standard input protocol in the User Guide for Animal Tissues (document part number A160162 v01) and starting from 127 mg of fresh frozen heart tissue. Final library quality was assayed as above in addition to qPCR using the Kapa Library quantification kit for Illumina (Roche Inc.). The indexed library was sequenced with other libraries on the Illumina NovaSeq SP flow cell with 2^*^150 bp paired end mode at the Norwegian Sequencing Centre.

### Genome assembly and curation, annotation and evaluation

A full list of relevant software tools and versions is presented in Supplementary Table 1. KMC (Kokot, Dlugosz, and Deorowicz 2017) was used to count k-mers of size 32 in the PacBio HiFi reads, excluding k-mers occurring more than 10,000 times. GenomeScope (Ranallo-Benavidez, Jaron, and Schatz 2020) was run on the k-mer histogram output from KMC to estimate genome size, heterozygosity and repetitiveness while ploidy level was investigated using Smudgeplot (Ranallo-Benavidez, Jaron, and Schatz 2020). HiFiAdapterFilt (Sim et al. 2022) was applied on the HiFi reads to remove possible remnant PacBio adapter sequences. The filtered HiFi reads were assembled using hifiasm (Cheng et al. 2021) with Hi-C integration resulting in a pair of haplotype-resolved assemblies, pseudo-haplotype one (hap1) and pseudo-haplotype two (hap2). Unique k-mers in each assembly/pseudo-haplotype were identified using meryl (Rhie et al. 2020) and used to create two sets of Hi-C reads, one without any k-mers occurring uniquely in hap1 and the other without k-mers occurring uniquely in hap2. K-mer filtered Hi-C reads were aligned to each scaffolded assembly using BWA-MEM (H. Li 2013) with -5SPM options. The alignments were sorted based on name using samtools (H. Li et al. 2009) before applying samtools fixmate to remove unmapped reads and secondary alignments and to add mate score, and samtools markdup to remove duplicates. The resulting BAM files were used to scaffold the two assemblies using YaHS (Zhou, McCarthy, and Durbin 2023) with default options. FCS-GX (Astashyn et al. 2024) was used to search for putative contamination. Contaminated sequences were removed. The mitochondrion was searched for in reads using Oatk (Zhou et al. 2024). Merqury (Rhie et al. 2020) was used to assess the completeness and quality of the genome assemblies by comparing to the k-mer content of the Hi-C reads. BUSCO (Manni et al. 2021) was used to assess the completeness of the genome assemblies by comparing against the expected gene content in the mammalia and cetartiodactyla lineages. Gfastats (Formenti et al. 2022) was used to output different assembly statistics of the assemblies. The assemblies were manually curated using PretextView and Rapid curation 2.0. Chromosomes (including sex chromosomes) were identified by mapping to GCA_949782905.1 (Svalbard reindeer) in addition to inspecting the Hi-C contact map in PretextView. BlobToolKit and BlobTools2 (Laetsch and Blaxter 2017), in addition to blobtk were used to visualize assembly statistics and GC-coverage plots. To generate the Hi-C contact map image, the Hi-C reads were mapped to the assemblies using BWA-MEM (H. Li 2013) using the same approach as above, before PretextMap was used to create a contact map which was visualized using PretextSnapshot.

**Table 1.**
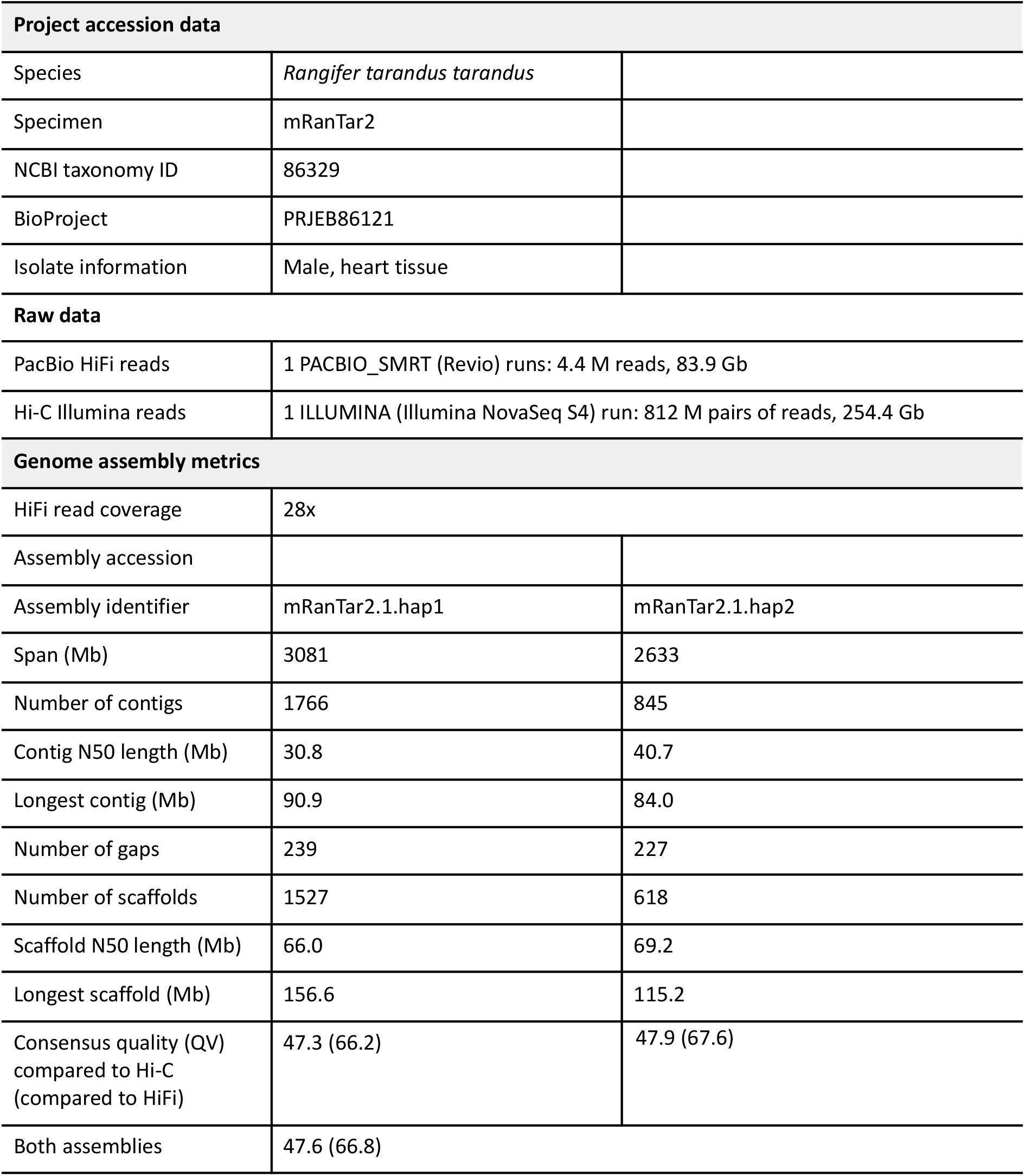
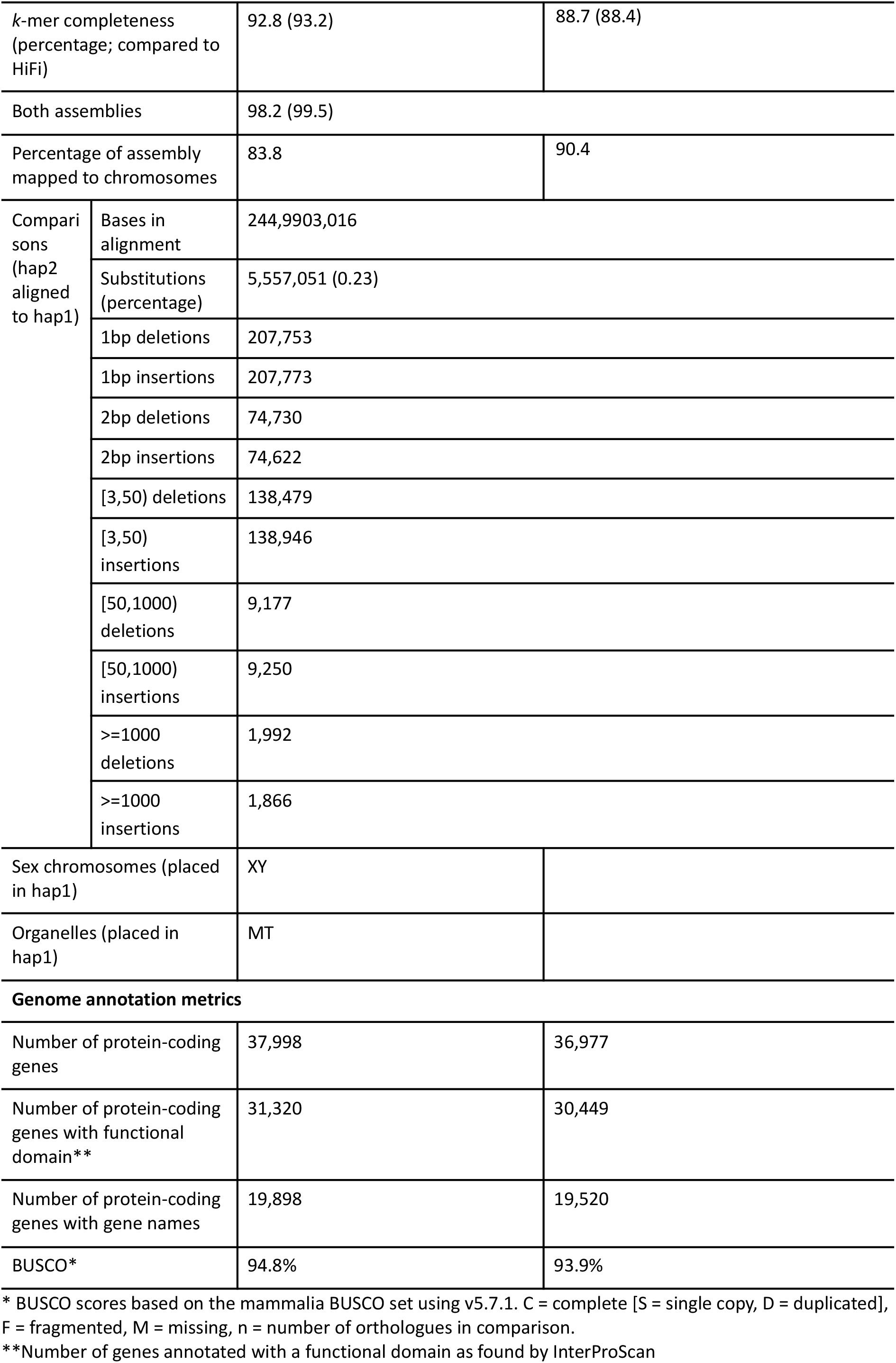
Genome data for *Rangifer tarandus tarandus*.

| Project accession data |  |  |
| --- | --- | --- |
| Species | *Rangifer tarandus tarandus* |  |
| Specimen | mRanTar2 |  |
| NCBI taxonomy ID | 86329 |  |
| BioProject | PRJEB86121 |  |
| Isolate information | Male, heart tissue |  |
| Raw data |  |  |
| PacBio HiFi reads | 1 PACBIO_SMRT (Revio) runs: 4.4 M reads, 83.9 Gb |  |
| Hi-C Illumina reads | 1 ILLUMINA (Illumina NovaSeq S4) run: 812 M pairs of reads, 254.4 Gb |  |
| Genome assembly metrics |  |  |
| HiFi read coverage | 28x |  |
| Assembly accession |  |  |
| Assembly identifier | mRanTar2.1.hap1 | mRanTar2.1.hap2 |
| Span (Mb) | 3081 | 2633 |
| Number of contigs | 1766 | 845 |
| Contig N50 length (Mb) | 30.8 | 40.7 |
| Longest contig (Mb) | 90.9 | 84.0 |
| Number of gaps | 239 | 227 |
| Number of scaffolds | 1527 | 618 |
| Scaffold N50 length (Mb) | 66.0 | 69.2 |
| Longest scaffold (Mb) | 156.6 | 115.2 |
| Consensus quality (QV) compared to Hi-C (compared to HiFi) | 47.3 (66.2) | 47.9 (67.6) |
| Both assemblies | 47.6 (66.8) |  |

|  |  |  |  |
| --- | --- | --- | --- |
| *k*-mer completeness (percentage; compared to HiFi) |  | 92.8 (93.2) | 88.7 (88.4) |
| Both assemblies |  | 98.2 (99.5) |  |
| Percentage of assembly mapped to chromosomes |  | 83.8 | 90.4 |
| Comparisons (hap2 aligned to hap1) | Bases in alignment | 244,9903,016 |  |
|  | Substitutions (percentage) | 5,557,051 (0.23) |  |
|  | 1bp deletions | 207,753 |  |
|  | 1bp insertions | 207,773 |  |
|  | 2bp deletions | 74,730 |  |
|  | 2bp insertions | 74,622 |  |
|  | [3,50) deletions | 138,479 |  |
|  | [3,50) insertions | 138,946 |  |
|  | [50,1000) deletions | 9,177 |  |
|  | [50,1000) insertions | 9,250 |  |
|  | >=1000 deletions | 1,992 |  |
|  | >=1000 insertions | 1,866 |  |
| Sex chromosomes (placed in hap1) |  | XY |  |
| Organelles (placed in hap1) |  | MT |  |
| **Genome annotation metrics** |  |  |  |
| Number of protein-coding genes |  | 37,998 | 36,977 |
| Number of protein-coding genes with functional domain** |  | 31,320 | 30,449 |
| Number of protein-coding genes with gene names |  | 19,898 | 19,520 |
| BUSCO* |  | 94.8% | 93.9% |

\* BUSCO scores based on the mammalia BUSCO set using v5.7.1. C = complete [S = single copy, D = duplicated], F = fragmented, M = missing, n = number of orthologues in comparison.
\*\*Number of genes annotated with a functional domain as found by InterProScan

We annotated the genome assemblies using a pre-release version of the EBP-Nor genome annotation pipeline (https://github.com/ebp-nor/GenomeAnnotation). First, AGAT (https://zenodo.org/record/7255559) agat_sp_keep_longest_isoform.pl and agat_sp_extract_sequences.pl were used on the GRCh38 genome assembly and annotation to generate one protein (the longest isoform) per gene. Miniprot (H. Li 2023) was used to align the proteins to the curated assemblies. UniProtKB/Swiss-Prot (UniProt Consortium 2023) release 2024_04 in addition to the vertebrata part of OrthoDB v11 (Kuznetsov et al. 2023) were also aligned separately to the assemblies. Red (Girgis 2015) was run via redmask (https://github.com/nextgenusfs/redmask) on the assemblies to mask repetitive areas. GALBA (Brůna et al. 2023; Buchfink, Xie, and Huson 2015; Hoff and Stanke 2019; H. Li 2023; Stanke et al. 2006) was run with the GRCh38 proteins using the miniprot mode on the masked assemblies. The funannotate-runEVM.py script from Funannotate was used to run EvidenceModeler (Haas et al. 2008) on the alignments of GRCh38 proteins, UniProtKB/Swiss-Prot proteins, vertebrata proteins and the predicted genes from GALBA. The resulting predicted proteins were compared to the protein repeats that Funannotate distributes using DIAMOND blastp and the predicted genes were filtered based on this comparison using AGAT. The filtered proteins were compared to the UniProtKB/Swiss-Prot release 2024_04 using DIAMOND (Buchfink, Xie, and Huson 2015) blastp to find gene names and InterProScan was used to discover functional domains. AGATs agat_sp_manage_functional_annotation.pl was used to attach the gene names and functional annotations to the predicted genes. EMBLmyGFF3 (Norling, Jareborg, and Dainat 2018) was used to combine the fasta files and GFF3 files into a EMBL format for submission to ENA.

To characterize the differences between the two pseudo-haplotypes, we ran minimap2 (H. Li 2017) on the two pseudo-haplotypes. The resulting alignment was processed with paftools.js (packaged with minimap2), producing a report listing the number of insertions, SNPs and indels between the two pseudo-haplotypes.

To compare the assemblies generated in this study to other available reindeer genome assemblies, we downloaded GCA_963422255.1, GCA_902712895.1, GCA_022457185.1, GCA_019903745.2, GCA_014898785.1 and GCA_004026565.1 from NCBI, generated assembly metrics with gfastats and completeness with BUSCO and the cetartiodactyla lineage.

The text in Methods and parts of Results is based on a template we use for all the species we publish in the EBP-Nor project.

## Results

### *De novo* genome assembly and annotation

The genome from a wild alpine male reindeer (Figure 1), had an estimated genome size of 2.42 Gbp, with 0.53 % heterozygosity and a bimodal distribution based on the k-mer spectrum (Supplementary Figure 1). The clear shoulder on the left signifies the k-mers corresponding to the heterozygous regions of the genome, while the main peak is from homozygous regions (Supplementary Figure 1). A total of 28-fold coverage in Pacific Biosciences single-molecule HiFi long reads and 85-fold coverage in Arima Hi-C reads resulted in two haplotype-separated assemblies. The final assemblies have total lengths of 3,081 Mb and 2,633 Mb (Table 1 and Figure 2), respectively. Both of these are slightly larger than the k-mer based estimation, a difference that might be due to k-mers occurring more than 10,000 times are being excluded. Pseudo-haplotypes one (hap1) and two (hap2) have scaffold N50 size of 66.0 Mb and 69.2 Mb, respectively, and contig N50 of 30.8 Mb and 40.7 Mb, respectively (Table 1 and Figure 2). 34 autosomes were identified in both pseudo-haplotypes (numbered the same as in GCA_949782905.1; Svalbard reindeer) and the X and Y chromosomes were added to pseudo-haplotype one.

**Figure 1.**
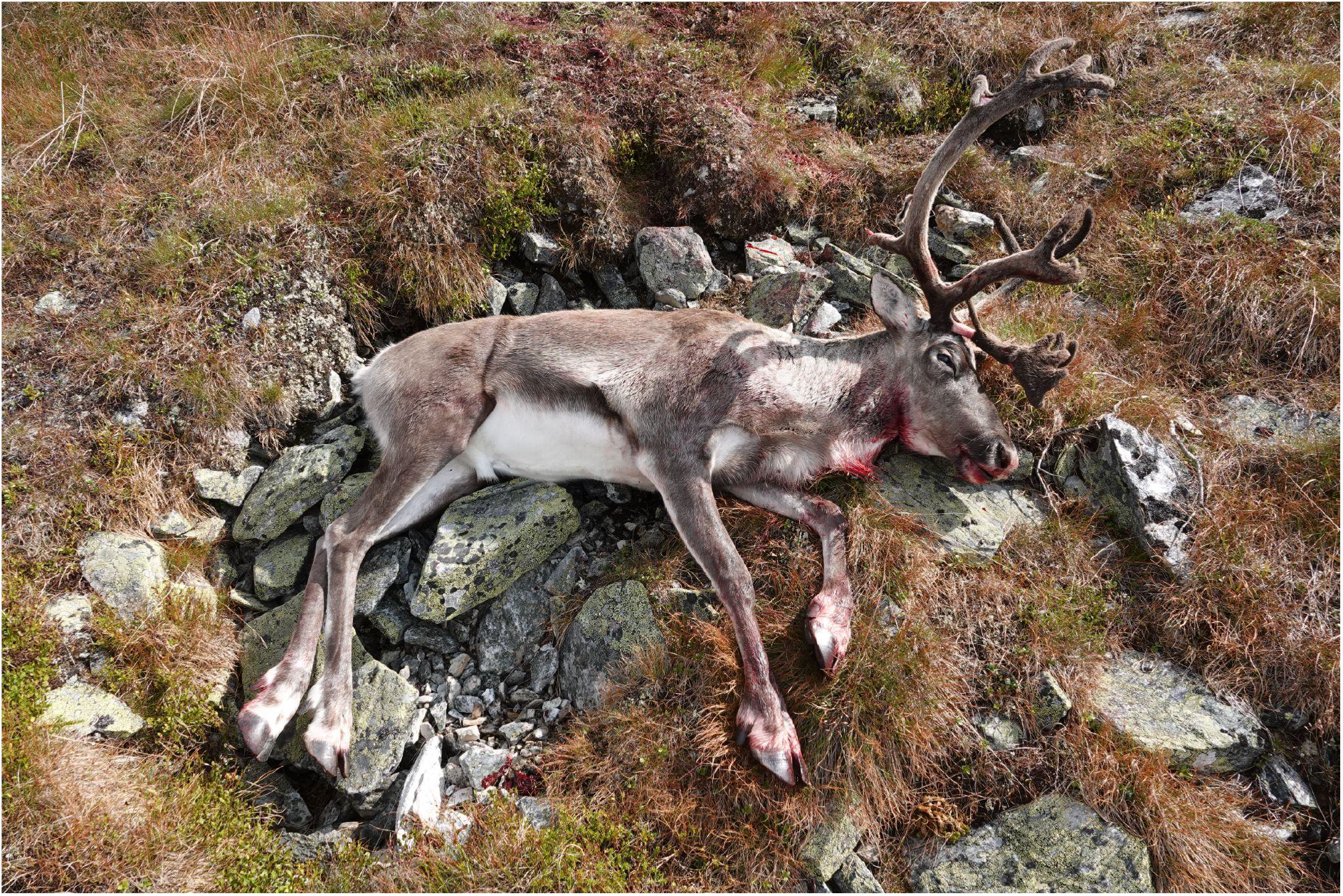
Sequenced specimen. Photograph of the wild alpine male specimen used for genome sequencing. The animal was legally shot during the ordinary hunting season on August 29, 2023 (Photo Anne Eline Streitlien).

**Figure 2.**
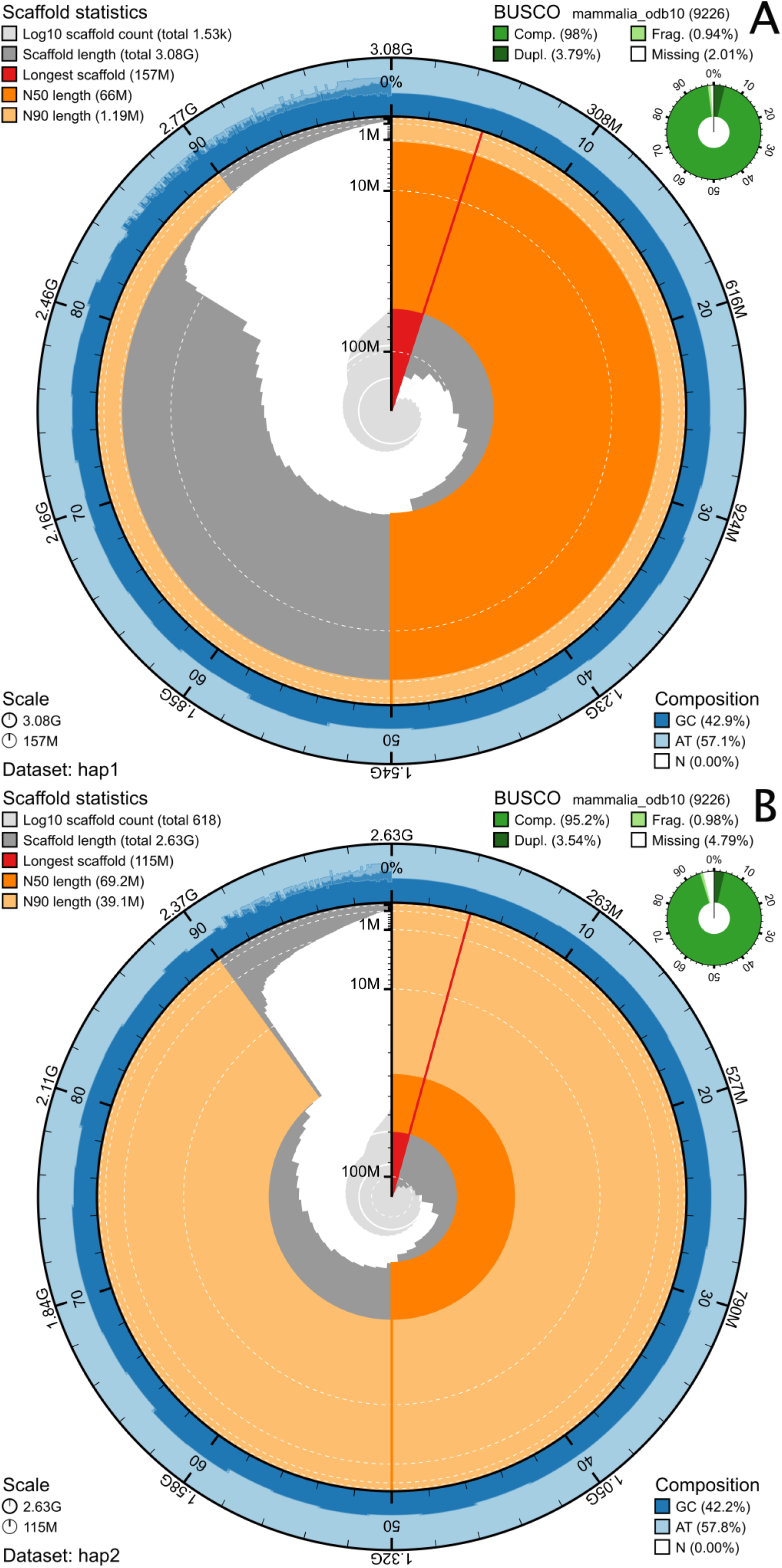
Metrics of the genome assemblies of *Rangifer tarandus tarandus*, A: hap1 and B: hap2. The BlobToolKit Snailplots show N50 metrics and BUSCO gene completeness. The two outermost bands of the circle signify GC versus AT composition at 0.1% intervals. Light orange shows the N90 scaffold length, while the deeper orange is N50 scaffold length. The red line shows the size of the largest scaffold. All the scaffolds are arranged in a clockwise manner from the largest to the smallest and are shown in darker gray with white lines at different orders of magnitude. The light gray shows the cumulative scaffold count. The scale inset in the lower left corner shows the total amount of sequence in the whole circle, and the fraction of the circle encompassed in the largest scaffold.

In addition to the alpine reindeer, we reassembled the previously published Svalbard reindeer (Dussex et al. 2023) due to changes and improvements in the pipeline used, resulting in scaffold N50 size of 70.7 Mb and 66.9 Mb, respectively, and contig N50 of 45.9 Mb and 48.1 Mb, in pseudo-haplotypes one and two respectively (Table 2).

**Table 2.** Assembly metrics for various reindeer genome assemblies.

| Assembly | Subspecies | Assembly length (Mb) | N50 contig (Mb) | N50 scaffold (Mb) | Chromosome level/sex chromosomes | Annotation available | Sequencing data |
| --- | --- | --- | --- | --- | --- | --- | --- |
| mRanTar2.1.hap1 | *R. t. tarandus* | 3,081 | 30.8 | 66.0 | Yes/Yes | Yes | PacBio HiFi |
| mRanTar2.1.hap2 | *R. t. tarandus* | 2,633 | 40.7 | 69.2 | Yes/No | Yes | PacBio HiFi |
| mRanTar1.2.hap1 | *R. t. platyrhynchus* | 2,990 | 45.9 | 70.7 | Yes/Yes | Yes | PacBio HiFi |
| mRanTar1.2.hap2 | *R. t. platyrhynchus* | 2,821 | 48.1 | 66.9 | Yes/No | Yes | PacBio HiFi |
| GCA_963422255.1 | *R. t. fennicus* (?) | 2,663 | 0.049 | 69.7 | No/No |  | Illumina + Hi-C |
| GCA_902712895.1 | *R. t. tarandus* | 2,490 | 0.196 | 20.7 | No/No |  | Illumina |
| GCA_022457185.1 | Not specified | 2,832 | 0.092 | 0.98 | No/No |  | Illumina |
| GCA_019903745.2 | *R. t. caribou* | 2,577 | 0.164 | 54.4 | Yes/No |  | Illumina, PacBio CLR and FISH |
| GCA_014898785.1 | *R. t. granti* | 2,537 | 0.231 | 9.35 | No/No |  | Illumina |
| GCA_004026565.1 | Not specified | 2,897 | 0.078 | 0.089 | No/No |  | Illumina |

Hap1 had 98.0% and hap2 95.2% complete BUSCO genes using the mammalian lineage set (Figure 2). When compared to a k-mer database of the Hi-C reads, hap1 had a k-mer completeness of 92.8%, pseudo-haplotype two of 88.7%, and combined they have a completeness of 98.2% (Supplementary Figure 2, also see Supplementary Figure 3). Further, hap1 has an assembly consensus quality value (QV) of 47.3 and pseudo-haplotype two of 47.9, where a QV of 40 corresponds to one error every 10,000 bp, or 99.99% accuracy compared to a k-mer database of the Hi-C reads (QV 66.2 and 67.9, respectively, compared to a k-mer database of the HiFi reads) (Table 1). The Hi-C contact map for the assemblies are shown in Supplementary Figure 4, and show clear separation of the different chromosomes. GC-coverage plots for the assemblies are found in Supplementary Figure 5, showing similar coverage in the chromosomes with some spread in GC content.

When comparing the two pseudo-haplotypes using minimap2, there are 5,557,051 SNPs differences (0.23 % of the aligned sequence), 422,954 deletions in hap2 compared to hap1 ranging from 1 bp to more than 1000 bp and 423,207 insertions from 1 bp to more than 1000 bp in size (Table 1). A total of 37,998 and 36,977 protein-coding genes were annotated in pseudo-haplotype one and two, respectively (Table 1).

The assemblies generated in this study all have chromosome-level scaffolds, while only one other reindeer genome assembly has this (GCA_019903745.2) (Table 2). With the exception of the Svalbard reindeer assembly, none of the other publicly available genome assemblies have N50 contig length of more than 1 Mb. All the downloaded assemblies have shorter assembly lengths than the around 3,000 Mb for hap1 for both the alpine reindeer and Svalbard reindeer, in some cases more than 500 Mb shorter (Table 2).

Most of the reindeer genome assemblies show good representation of complete BUSCO genes, with only one (GCA_004026565.1) having fewer than 90 % complete genes (Table 3).

**Table 3.**
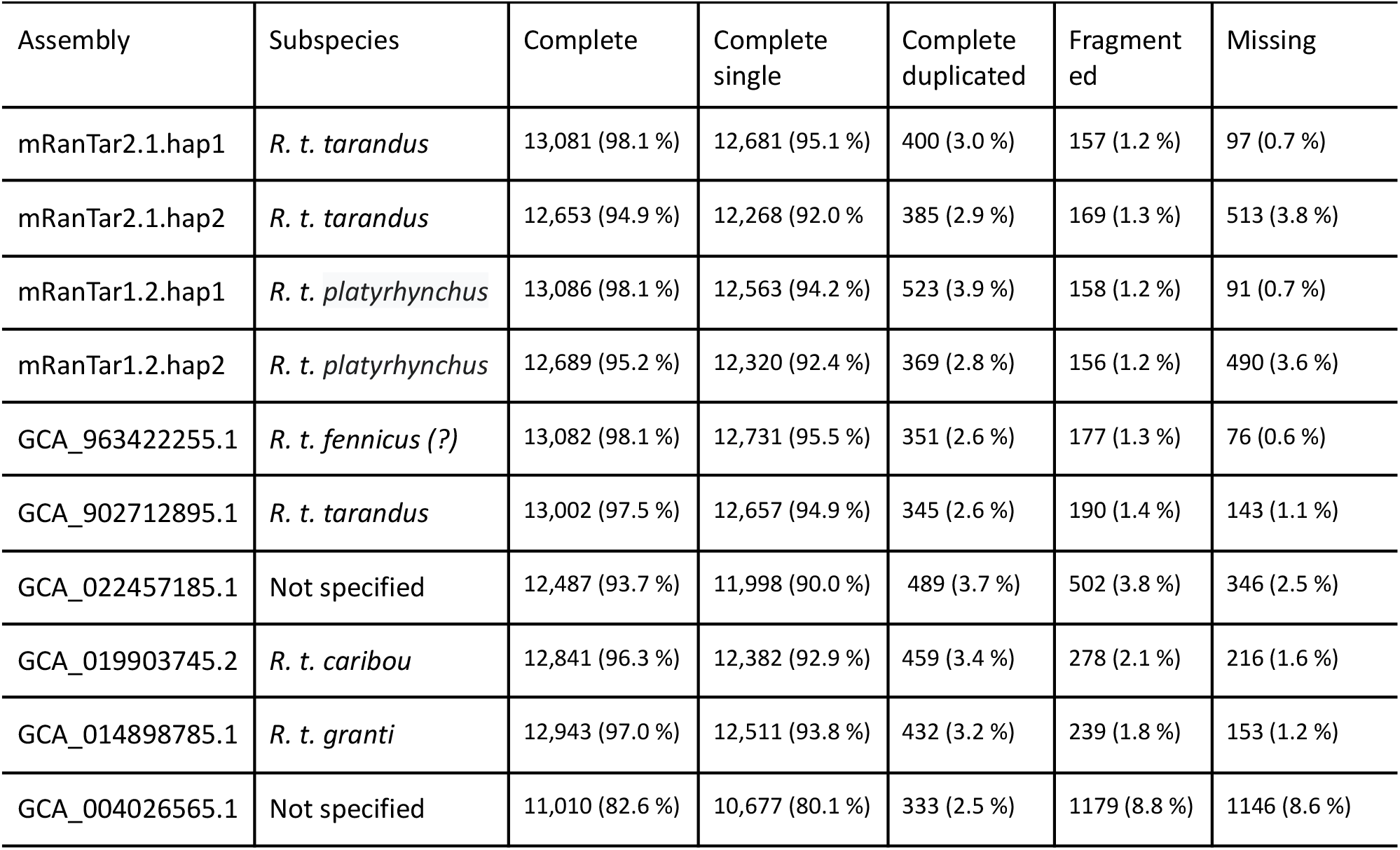
BUSCO scores for various reindeer genome assemblies.

## Discussion

We have sequenced, assembled, and annotated chromosome-level, pseudo-haplotype separated genome assemblies from *R. tarandus tarandus* from the Rondane area in Norway. The Rondane population represents the native wild reindeer (Kvie et al. 2019). This is the second individual sequenced with long-read data, with the Svalbard reindeer done earlier (Dussex et al. 2023). For the Svalbard reindeer, we have reassembled the sequencing data previously generated and annotated the resulting pseudo-haplotype separated assembly, resulting in higher contig N50 lengths (45.9 and 48.1 Mb in the new version compared to 22.5 and 25.5 Mb in the original; Table 2), as well as better completeness with regards to BUSCO genes (98.1 % and 95.2 compared to 96.3 % and 94.1 %; Table 3) (Dussex et al. 2023).

There are currently 8 genome assemblies for reindeer available in INSDC (i.e., ENA, NCBI and DDBJ), including the two pseudo-haplotypes for Svalbard reindeer we published earlier (Dussex et al. 2023). Some of these are generated from individuals from specific sub-species, while this is not specified for others (Table 2 and 3). Which genome assembly to use as a reference for a specific comparative or population genomic study will depend on the particulars of that study. For instance, even though the different assemblies vary substantially in contiguity (from 0.049 to 48.1 Mb N50 contig, almost thousandfold difference), the variation in complete BUSCO genes is much less (82.6 to 98.1 %), illustrating that it is easier to assemble unique regions (e.g., genes) of a genome than the repetitive regions.

The previous Svalbard reindeer genome assembly (updated in this study) and this wild alpine reindeer genome are the only two of the available assemblies providing genome annotation at INSDC. While there might be some annotations available for some of the other genomes, these are hard to locate since they are not linked from INSDC. The two genome assemblies presented here (wild alpine and Svalbard) are substantially more contiguous than other reindeer assemblies, have high amounts of complete BUSCO genes and available annotations and will serve as excellent references for population genomic analyses. These annotated genomes will be valuable resources for studies addressing genes under selection in various scenarios such as between wild and domesticated reindeer and will facilitate further genomic research on ancestry, genetic diversity, inbreeding, genetic drift, population structure, domestication, feralization and local adaptation of the reindeer, which may hopefully contribute towards preserving the last populations of wild alpine reindeer.

## Acknowledgements

This project received data management and infrastructure support from ELIXIR Norway, supported by the Research Council of Norway’s grant 270068, the University of Bergen, the University of Oslo, the Arctic University of Norway in Tromsø, the Norwegian University of Science and Technology and the Norwegian University of Life Sciences: NMBU. The authors acknowledge support from the National Infrastructure for High Performance Computing and resources provided by Sigma2 as well as Data Storage in Norway (project NN8013K) for computational work. The Norwegian Sequencing Centre generated the sequencing data used in this project (http://sequencing.uio.no).

## Authors’ contributions’

**Ole K. Tørresen:** Writing - original draft, Formal analysis, Visualization, Writing - review and editing. **Anne Eline Streitlien:** Resources. **Morten Skage:** Investigation. **Giada Ferrari:** Investigation. **Ave Tooming-Klunderud:** Investigation, Writing - original draft. **Jose Cerca:** Investigation. **Christer M. Rolandsen:** Resources. **Olav Strand:** Resources. **Atle Mysterud:** Funding acquisition, Project administration, Writing - original draft, Writing - review and editing. **Kjetill S. Jakobsen:** Funding acquisition, Project administration, Writing - original draft, Writing - review and editing,

## Data Availability

The gene annotations and genome assemblies are available at 10.5281/zenodo.15342678.

