## Supplementary information for "Chromosome-level genome assembly of Norwegian wild alpine reindeer (*Rangifer tarandus tarandus*)"

### Supplementary Material

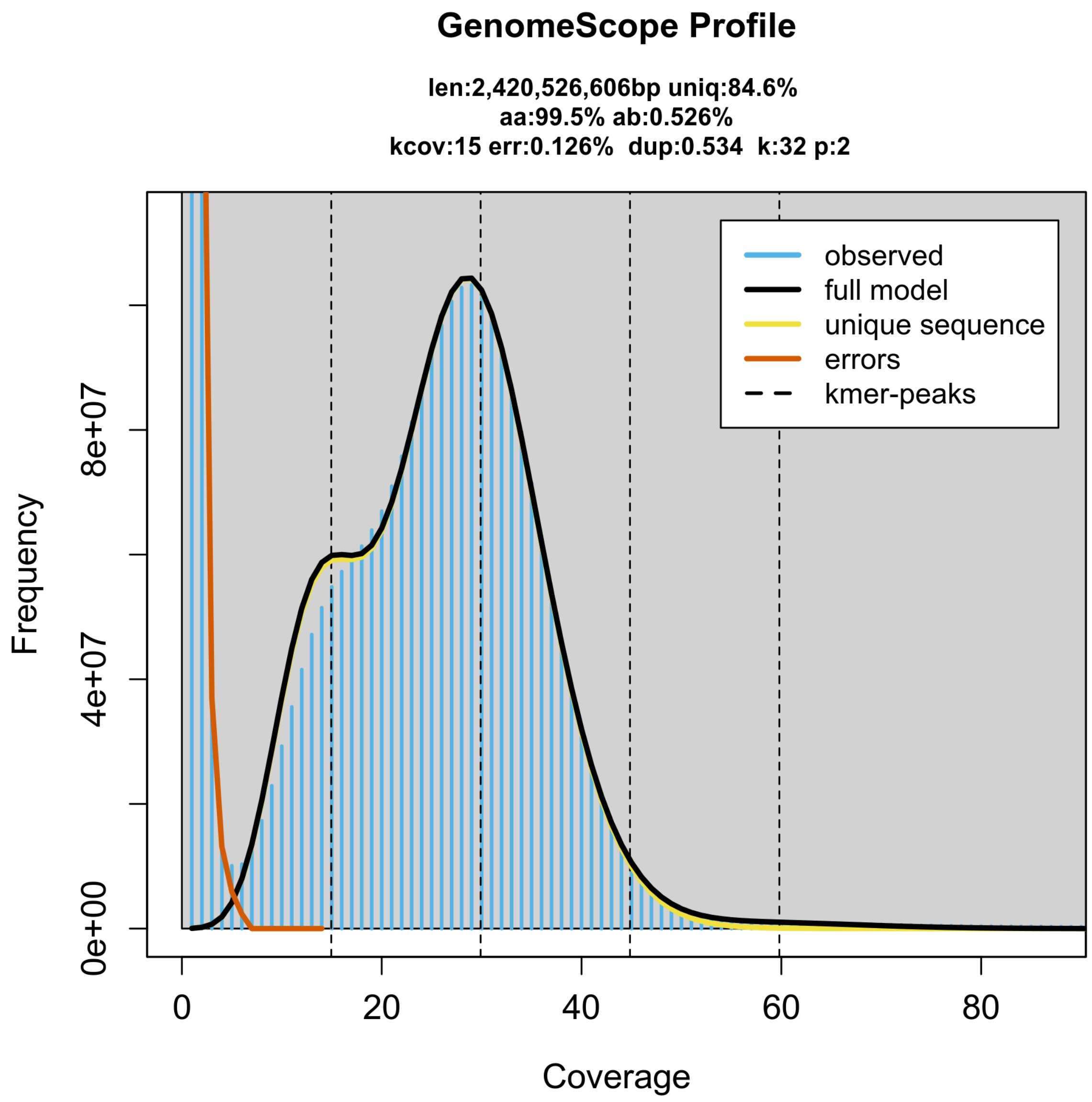


**Supplementary Figure 1: GenomeScope profile of the HiFi reads from the sequenced individual.** This analysis estimates a 2,420 Mb genome, with 0.53 % heterozygosity.

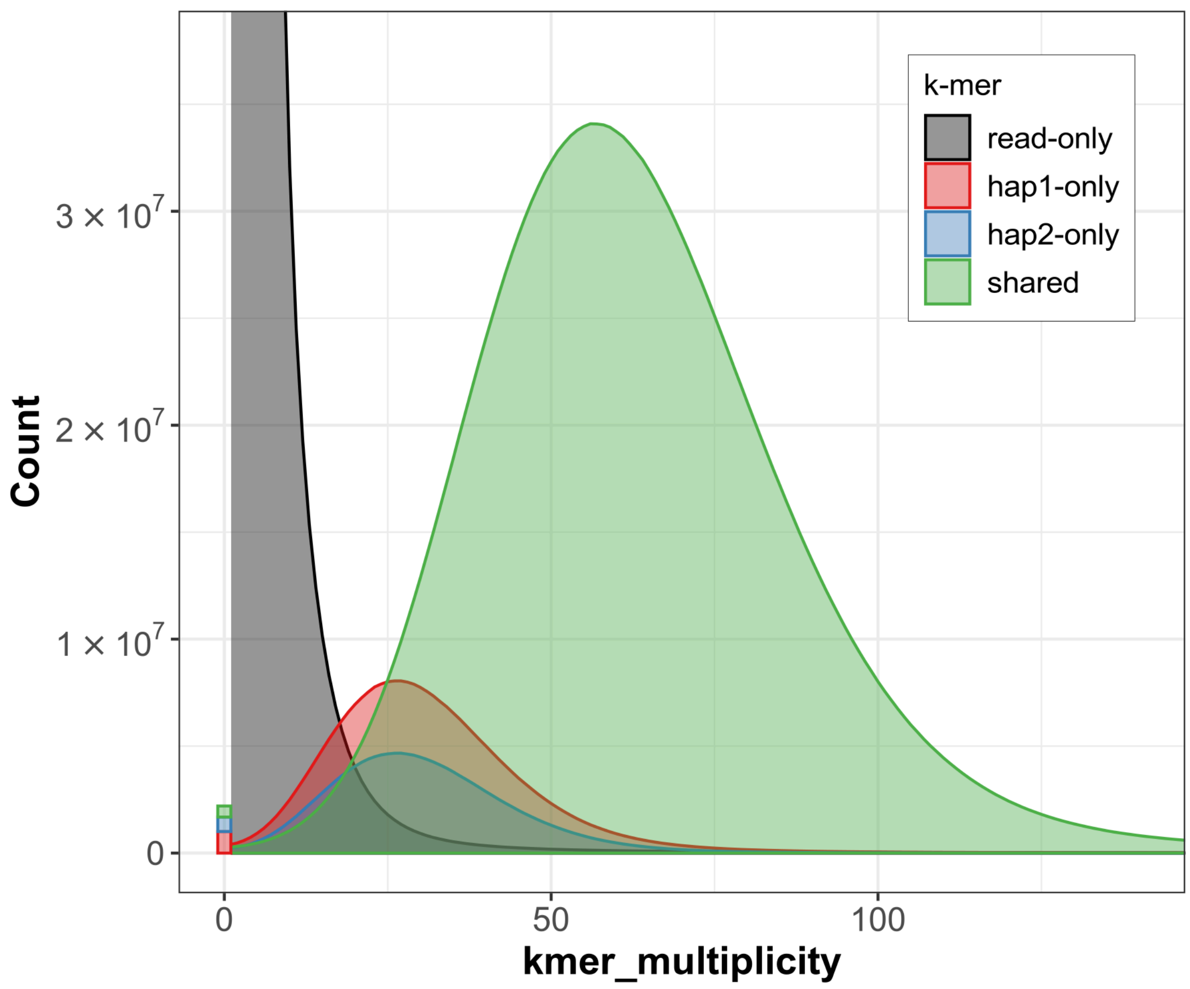


**Supplementary Figure 2: K-mer copy-number spectrum analysis of *Rangifer tarandus tarandus* compared to k-mers from a database from the Hi-C reads.** Assembly-specific k-mers are shown in red and blue, while k-mers shared by both pseudo-haplotypes are in green. The stack above 0 on the x-axis shows k-mers found in the assemblies, but not in the reads. Figure is generated by Merqury.

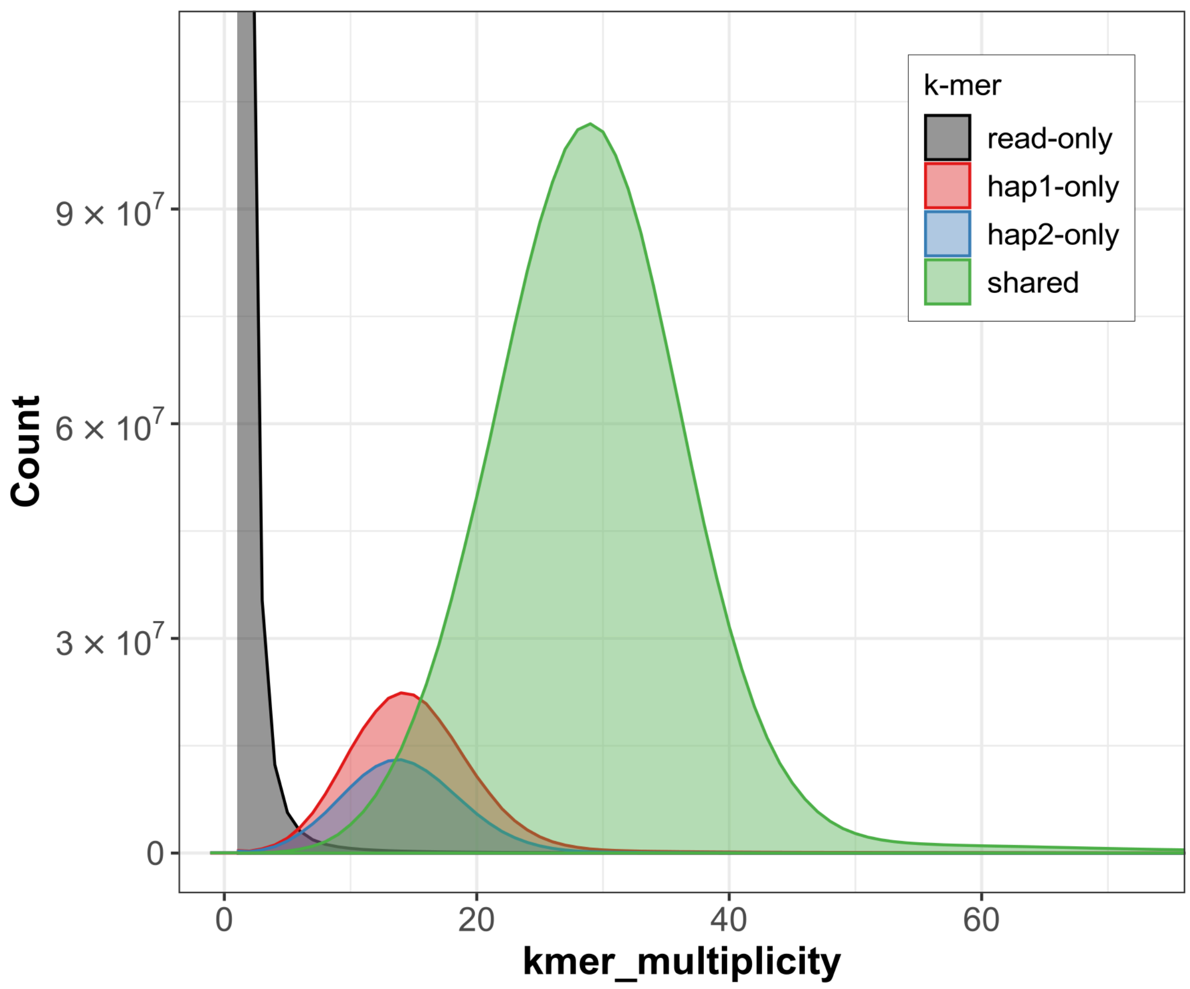


**Supplementary Figure 3: K-mer copy-number spectrum analysis of *Rangifer tarandus tarandus* compared to k-mers from a database from the HiFi reads.** Assembly-specific k-mers are shown in red and blue, while k-mers shared by both pseudo-haplotypes are in green. The stack above 0 on the x-axis shows k-mers found in the assemblies, but not in the reads. Figure is generated by Merqury.

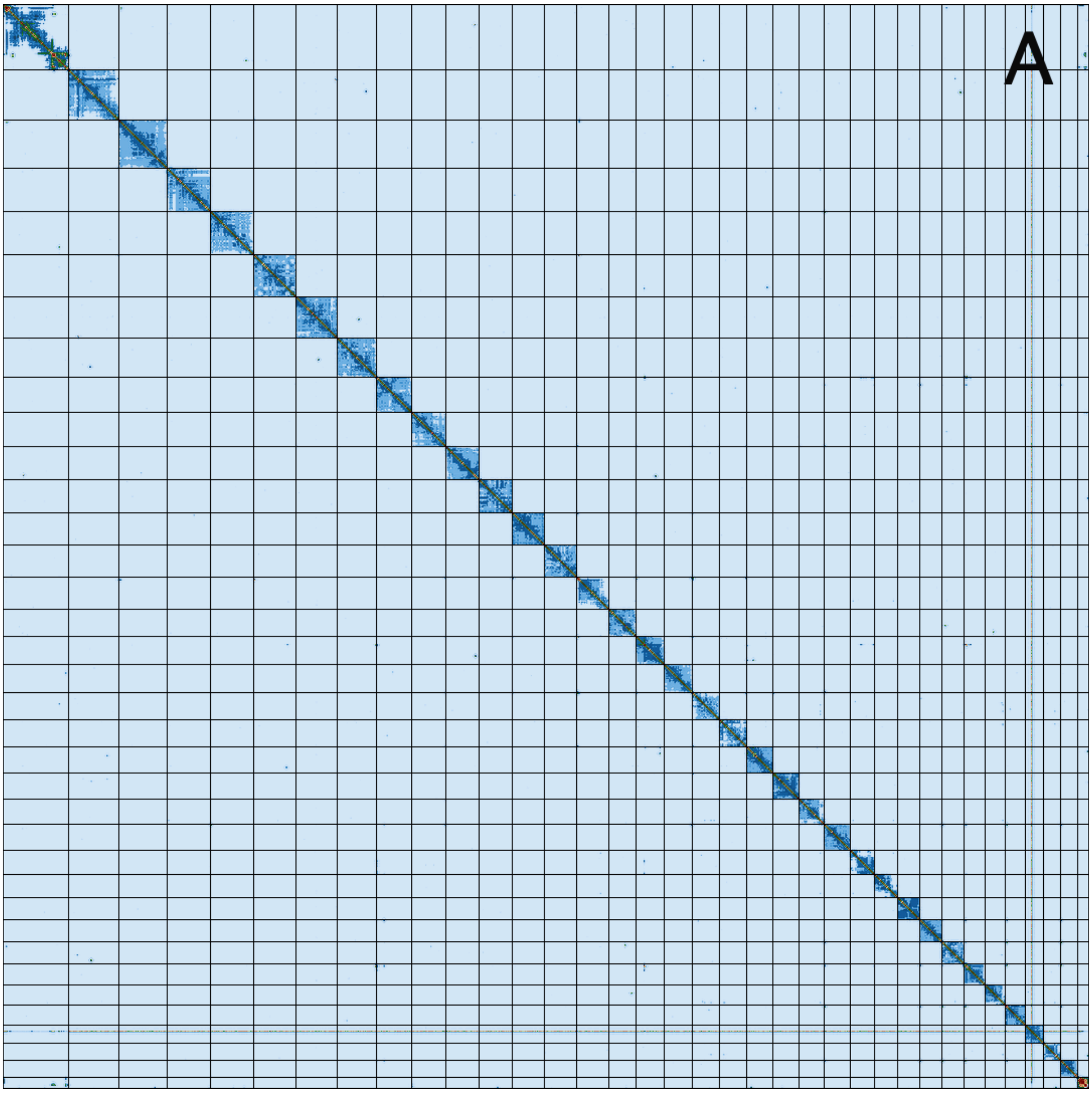
A

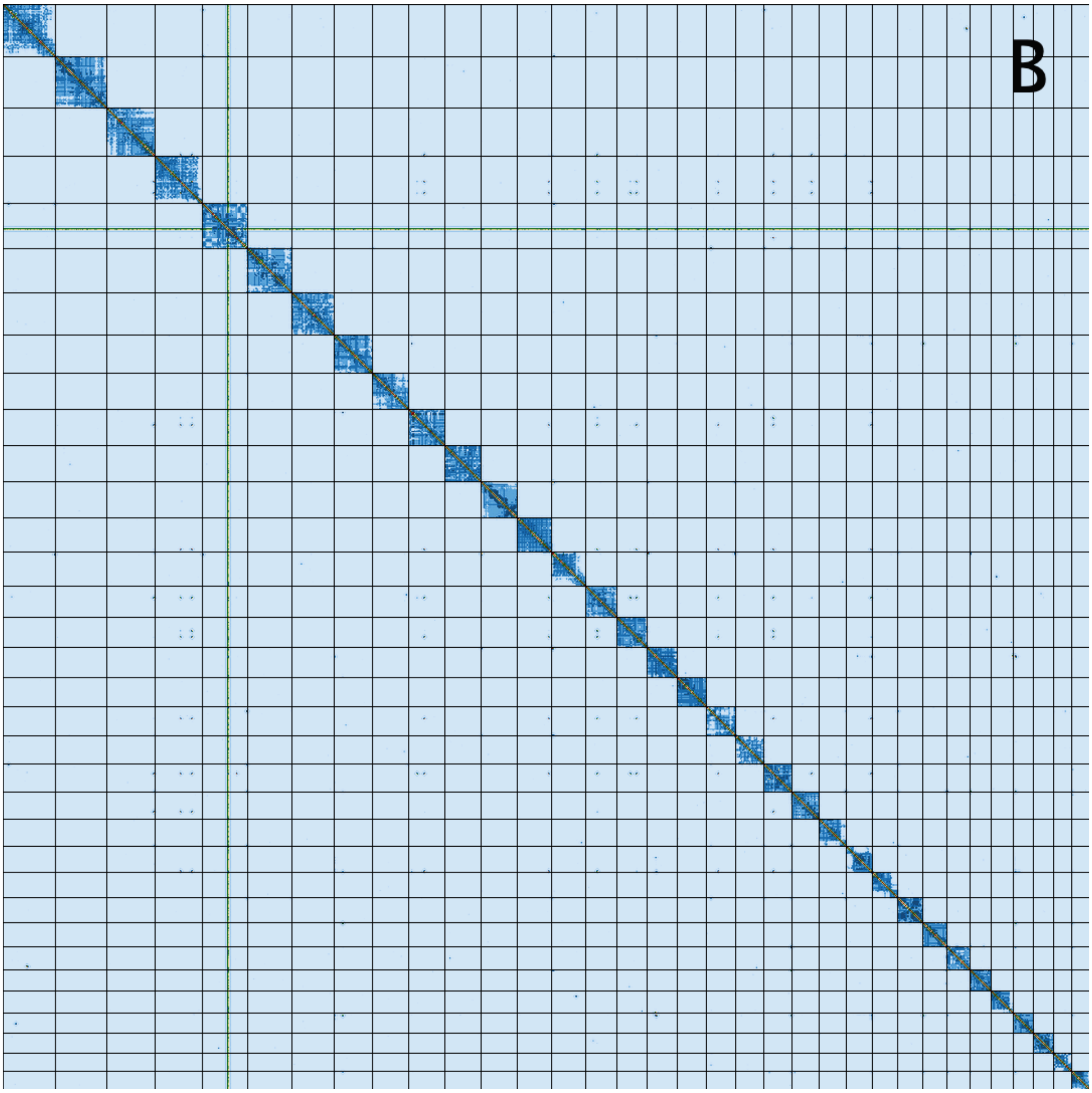


**Supplementary Figure 4: Hi-C contact map of genome assemblies of *R. t. tarandus* for hap1 (A) and hap2 (B).** The assemblies are visualized using PreTextSnapshot. Chromosomes are shown in order of size from left to right and top to bottom.

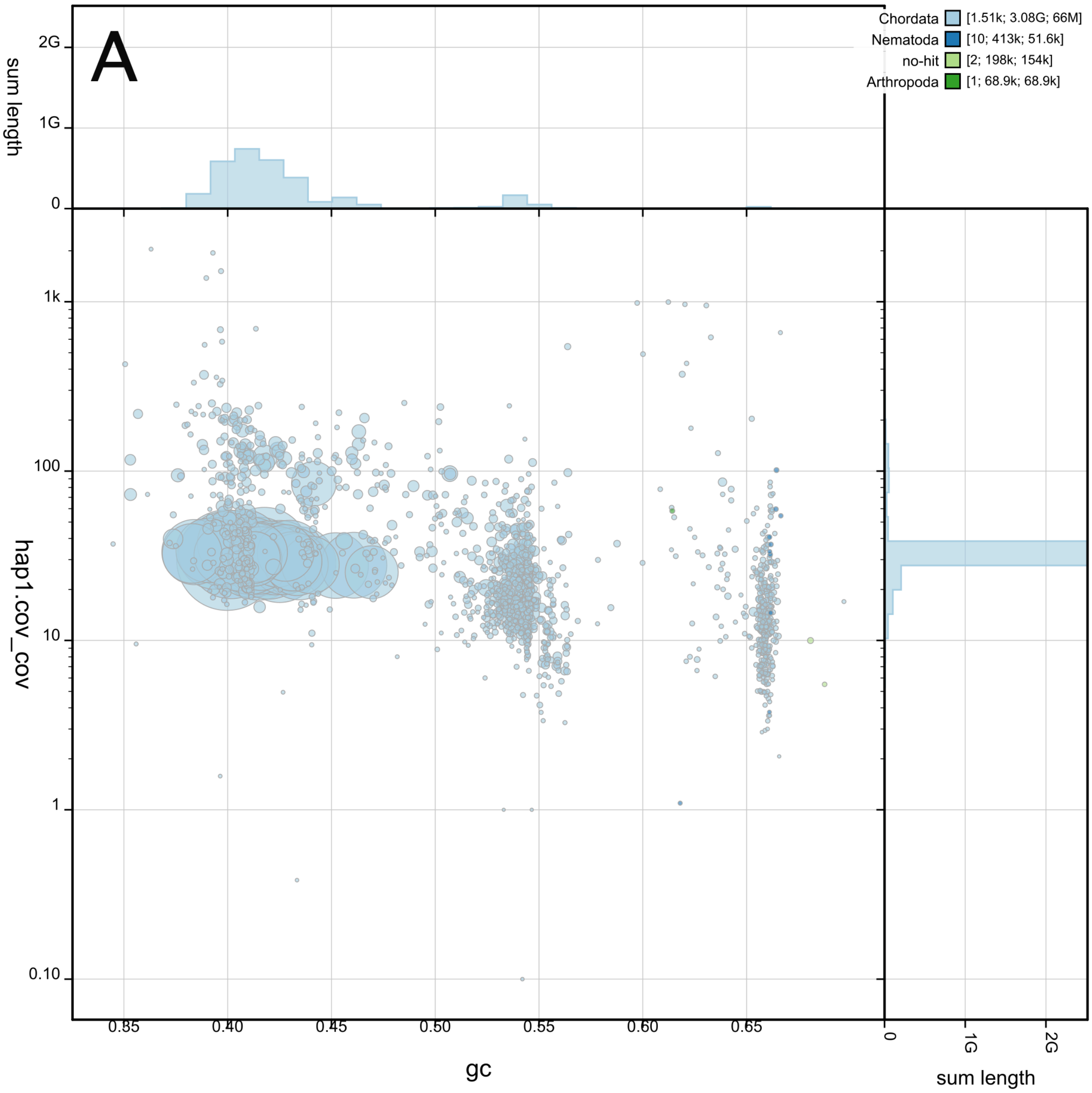
A
Chordata [1.51k; 3.08G; 66M]
Nematoda [10; 413k; 51.6k]
no-hit [2; 198k; 154k]
Arthropoda [1; 68.9k; 68.9k]
sum length
2G
1G
0
hap1.cov_cov
1k
100
10
1
0.10
0.35
0.40
0.45
0.50
0.55
0.60
0.65
gc
0
1G
2G
sum length

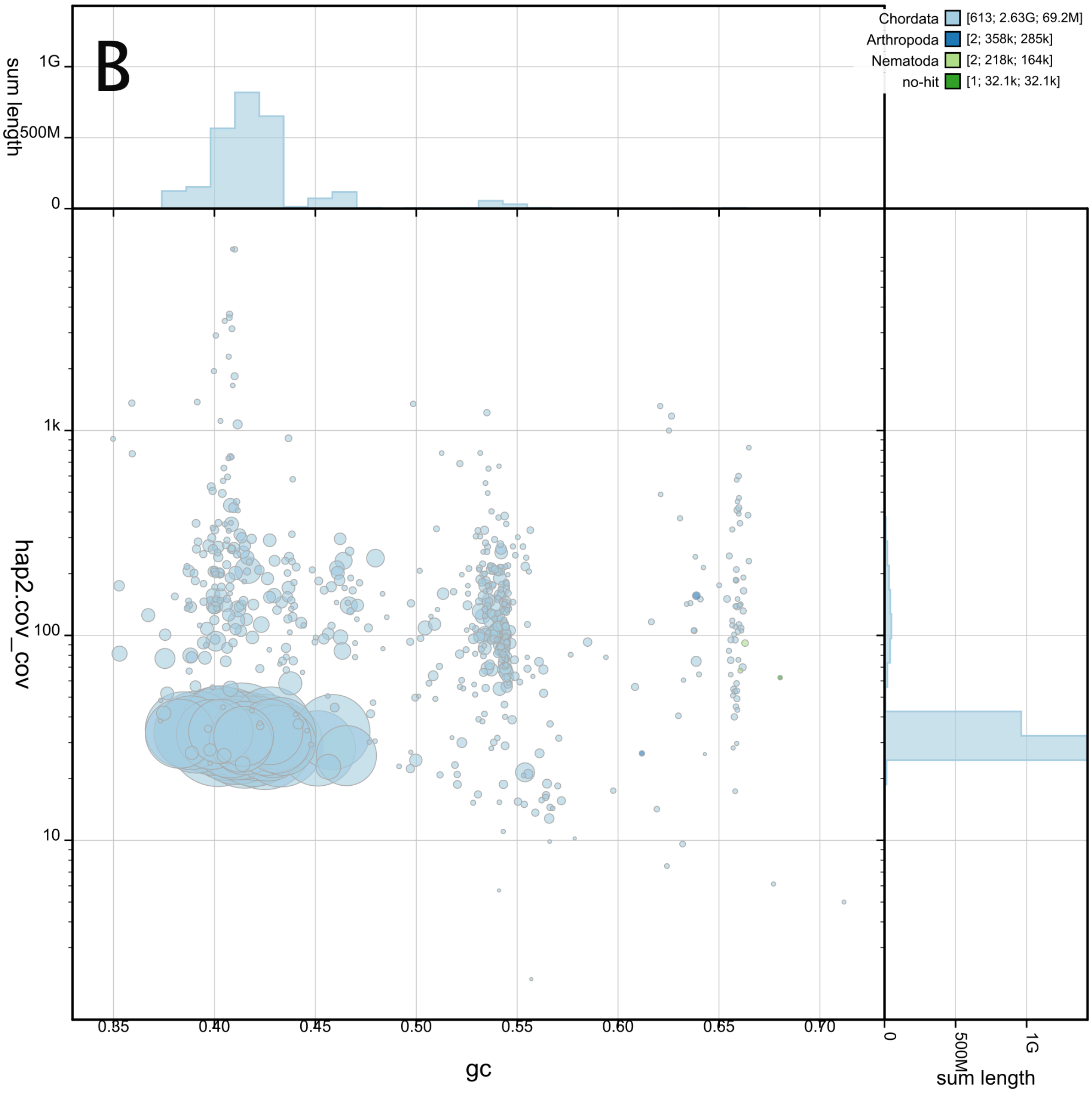


**Supplementary Figure 5: BlobToolKit GC-coverage plots of genome assemblies of *Rangifer tarandus tarandus* hap1 (A) and hap2 (B).** The scaffolds are coloured by phylum. The size of the circles are in proportion to the length of the scaffolds. Histograms show the distribution of scaffold length sum along each axis.

**Supplementary Table 1. Software tools:** versions and sources

| Software tool | Version | Source |
|---|---|---|
| BlobToolKit | 4.1.7 | https://github.com/blobtoolkit/blobtoolkit |
| blobtk | 0.5.8 | https://github.com/blobtoolkit/blobtk |

| | | |
|---|---|---|
| BUSCO | 5.7.1 | https://gitlab.com/ezlab/busco |
| hifiasm | 0.19.8 | https://github.com/chhylp123/hifiasm |
| KMC | 3.1.2 | https://github.com/refresh-bio/KMC |
| GenomeScope | 2.0 | https://github.com/tbenavi1/genomescope2.0 |
| HiFiAdapterFilt | 2.0.0 | https://github.com/sheinasim/HiFiAdapterFilt |
| PretextView | 0.2.5 | https://github.com/wtsi-hpag/PretextView |
| PretextMap | 0.1.9 | https://github.com/wtsi-hpag/PretextMap |
| PretextSnapshot | | https://github.com/wtsi-hpag/PretextSnapshot |
| meryl | 1.3.0 | https://github.com/marbl/meryl |
| BWA-MEM | 0.7.17 | https://github.com/lh3/bwa |
| samtools | 1.17 | https://github.com/samtools/samtools |
| YaHS | 1.2a.2 | https://github.com/c-zhou/yahs |
| FCS-GX | 0.4.0 | https://github.com/ncbi/fcs |
| Merqury | 1.3 | https://github.com/marbl/merqury |
| AGAT | 1.0 | https://github.com/NBISweden/AGAT |
| MitoHiFi | 2.2 | https://github.com/marcelauliano/MitoHiFi |
| miniprot | 0.13 | https://github.com/lh3/miniprot |
| GALBA | 1.0.6 | https://github.com/Gaius-Augustus/GALBA |
| RED | 2018.09.10 | http://toolsmith.ens.utulsa.edu/ |
| Funannotate | 1.8.17 | https://github.com/nextgenusfs/funannotate |

| | | |
|---|---|---|
| EvidenceModeler | 2.1.0 | https://github.com/EVidenceModeler/EVidenceModeler |
| DIAMOND | 2.1.8 | https://github.com/bbuchfink/diamond |
| InterProScan | 5.62-94 | https://www.ebi.ac.uk/interpro/search/sequence/ |
| EMBLmyGFF3 | 2.2 | https://github.com/NBISweden/EMBLmyGFF3 |
| Rapid curation 2.0 | 964d17e997e00c69f25940cf96d3658bda631147 | https://github.com/Nadolina/Rapid-curation-2.0, |
